# Neural Representations of Embodied Musical Emotions

**DOI:** 10.64898/2026.09.16.751954

**Authors:** Maya Rassouli, Viljami Salmela, Kerttu Seppälä, Harri Harju, Lauri Nummenmaa, Vesa Putkinen

## Abstract

Bodily sensations shape the emotional experience of music, yet the neural underpinnings of embodied musical emotions remain poorly understood. Here we investigated the correspondence between neural activity patterns and self-reported bodily sensations induced by emotional music. Participants (n=74) listened to emotionally engaging (aggressive, happy, sad, scary) music while their hemodynamic brain activity was measured with functional magnetic resonance imaging (fMRI). Music-induced bodily sensations were measured after imaging. Representational similarity analysis (RSA) was used to identify brain regions where the representational geometries of neural responses to music align with the topographical patterns of the corresponding bodily sensations, emotions, and musical features. Bodily sensations maps (BSMs) showed emotion-specific topographies. Univariate general linear model (GLM) analysis revealed increased neural activity within motor, somatosensory, interoceptive, limbic, and auditory brain regions during music listening, with little difference between the emotion categories. In contrast, RSA revealed significant correspondence between the representational geometries of neural activity patterns and emotion category-specific bodily sensations in motor, somatosensory, emotion-related and auditory brain regions. Multivariate pattern (MVP) classification analysis showed that activity in the auditory cortex accurately predicted the emotion categories. Our results indicate neural correspondence to bodily sensations in somatomotor and somatosensory brain regions, providing novel evidence for the neural basis of embodied music-induced emotions.

## Introduction

Music evokes strong bodily sensations, ranging from pleasurable chills to spontaneous movements. Such bodily responses are closely linked with how listeners perceive and emotionally engage with music (Burger et al., 2013). This relationship is central to theories of embodiment, which emphasize the interaction between bodily states, emotions, perception, and cognition (Nummenmaa et al., 2014, Leman & Maes, 2015), and distinct music-evoked emotions are associated with discernible patterns of bodily sensations (Putkinen et al., 2024). The subjective somatic sensations have been suggested to arise from the activation of the skeletal muscles along with the responses of the autonomic nervous system (ANS), and might play a role in the intensity of emotional responses as well as in distinguishing between different emotions such as joy, sadness, fear, aggressiveness, and tenderness (Putkinen et al., 2024). Neuroimaging studies have shown that emotional music engages auditory, limbic and paralimbic regions, but also somatosensory and motor cortices, indicating embodied representations of musical emotions (Putkinen et al., 2021; Koelsch et al., 2018; Koelsch et al., 2014). Furthermore, distinct emotions can be decoded from the neural activity patterns in the somatosensory and motor cortices (Putkinen et al., 2021), indicative of emotion-specific bodily representations of musical emotions.

While neural and bodily correlates of music-induced emotions have been mapped in neuroimaging and behavioural studies, the interaction between the brain and the body in these experiences remains poorly understood. Although traditional univariate analysis and multivariate pattern classification of neuroimaging data have demonstrated somatomotor involvement in music-induced emotions, these approaches do not reveal whether neural representations of musical emotions are organized according to subjective bodily experience, as predicted by theories of embodiment. It remains unknown whether the emotion-dependent somatosensory and motor activation patterns align with the actual bodily responses evoked by the music.

More broadly, music is a powerful tool for elucidating brain-body interaction in emotion given its unique ability to evoke strong emotions and subjective bodily experiences while stimulating several brain processes. Understanding music-induced brain-body processes may yield clinical value, as music has been increasingly employed in therapeutic and rehabilitative care of cognitive, emotional and motor impairments in e.g. stroke and Parkinson’s disease patients (Sihvonen et al., 2017; Forsblom et al., 2009, Pacchetti et al., 2000).

### The Current Study

Here, we examined the similarity between the representational geometry of the neural patterns and the self-reported bodily sensations associated with distinct music-induced emotions using functional magnetic resonance imaging (fMRI), emBODY tool (Nummenmaa et al., 2014) and representational similarity analysis (RSA) (Kriegeskorte et al., 2008). RSA enables comparison of the representational structure of neural activity patterns and subjective bodily sensation maps. This enables us to test whether relationships among music-induced bodily sensations are mirrored in distributed patterns of brain activity, thereby providing a direct link between subjective bodily experiences and neural representations. RSA was additionally performed using self-reported emotions as well as the musical features associated with the music excerpts. We hypothesized that the information about emotional embodiment during music listening is encoded in brain regions associated with motor control, auditory processing, somatosensation, interoception and emotional processing. Consistent with these hypotheses, the representational organization of music-evoked bodily sensations and that of neural responses corresponded in motor, somatosensory and auditory cortices, as well as in limbic and interoceptive brain regions.

## Materials and Methods

### Participants

74 volunteers (55 females, 19 males, mean age=28.84±9) participated in the study. Exclusion criteria were a past history of psychiatric or neurological disorders, alcohol or substance abuse, current use of medications affecting the central nervous system (e.g. SSRIs), and the standard MRI exclusion criteria assessed through self-report. All participants provided written informed consent and received compensation for their participation.

### Study Design

The participants underwent functional magnetic resonance imaging (fMRI) while listening to aggressive, happy, sad and scary music excerpts. These emotion categories were selected as they represent some of the most prominent emotions evoked by music and cover the four quadrants of the valence-arousal circumplex model (Eerola et al., 2010). The stimuli consisted of five 45-second music excerpts for each emotion category (20 in total), presented in a fixed pseudo-random order, with 15-second silent breaks between excerpts. The music excerpts were selected from Western popular music and movie soundtracks (**Supplementary Material Table S1. for full list**.) and were previously validated to elicit the intended emotional states and distinct bodily sensations (Putkinen et al., 2024). Binaural auditory stimuli were presented using MRI-compatible, individually adjusted headphones (Sensimetrics S14). Participants were instructed to focus on the music and remain still during the scan. High-resolution T1-weighted anatomical brain images were acquired at the beginning of the scan. After the scanning session, participants relistened to each musical clip and rated them for happiness, sadness, tenderness, fear, relaxation, energy, aggressiveness, danceability, liking, and irritation on a scale from 1 to 7 and reported their bodily sensations to each excerpt using the emBODY tool (Nummenmaa et al., 2014) in which the participants colored regions of a silhouette of a human body to indicate where they experienced activation.

### Statistical Analysis of Bodily Sensations, Emotion Ratings and Musical Features

To map where the different emotional categories were felt in the body, bodily sensation map (BSM) data for each emotion category were tested against zero using pixelwise one-sample t-tests across participants. In the resulting maps, pixel intensities represent the t-values for bodily changes associated with each music-induced emotion category across participants, thresholded using FDR correction at p < 0.05. For the subsequent RSA, uncorrected t-values were obtained similarly for each song.

Additionally, 20 musical features were extracted from the songs using the MIR toolbox version 1.7. (Lartillot & Toiviainen, 2007) in Matlab (see list of features in **Supplementary Material Table S2**.). A principal component analysis (PCA) was run to reduce the dimensionality of musical features into 3 principal components (PCs), and correlations between the song-wise mean emotion ratings and musical features component scores were calculated.

### fMRI Data Acquisition

A 3T GE SIGNA PET/MR system (General Electric Medical Systems, Milwaukee, Wisconsin) with a 48-channel head coil was used to collect the functional and structural MRI data. High-resolution anatomical T1-weighted (T1w) images were acquired with the Ultrafast Gradient Echo BRAVO sequence for anatomical normalization (1 mm3 resolution, TR 8.5 ms, TE 3.7 ms, flip angle 10°, 256 mm FOV, 256 x 256 reconstruction matrix). A total of 405 functional volumes (including 5 dummy volumes) were acquired with a T2*-weighted echo-planar imaging sequence sensitive to the blood-oxygen-level-dependent (BOLD) signal contrast (TR 3000 ms, TE 30 ms, 90° flip angle, 256 mm FOV, 96 × 96 reconstruction matrix, 2.7 mm slice thickness, 0 mm spacing, 51 interleaved axial slices acquired in ascending order.

### MRI Data Preprocessing

fMRIPrep 21.0.0rc.2 was used to preprocess both structural and functional MRI data. The anatomical T1w reference image preprocessing included intensity normalization to ensure comparability of intensity values across images, skull-tripping to remove non-brain tissue, brain tissue segmentation to classify different tissue types, spatial normalization to align anatomical images to MNI152NLin2009cAsym standard space and brain surface reconstruction using Freesurfer. Functional data preprocessing included functional-to-anatomical registration, slice timing correction, spatial smoothing with a Gaussian filter, motion correction using ICA-AROMA, and spatial normalization to MNI152NLin2009cAsym standard space.

### Full Volume General Linear Model Analysis

To identify brain regions activated by each music-induced emotion category as compared to periods of silence, a full-volume general linear model (GLM) was fitted using SPM12 in MATLAB. Each musical emotion category was modelled with separate boxcar regressors. Contrast images comparing each emotion category against baseline (silence) were generated for each subject. These images were then subjected to second-level analysis for population level inference. Any clusters that withstood false discovery rate (FDR) correction (*p* < 0.001 at voxel-level) are reported. For subsequent RSA (see below), a separate first-level model was fitted in which each song was modelled with individual regressors yielding contrast images reflecting the response to each song versus baseline (silence).

### Regions-of-Interest

For subsequent regional analyses, beta weights were extracted from 15 regions-of-interest (ROI): precentral gyrus (PRECG), postcentral gyrus (POSCG), supplementary motor area (SMA), cerebellum, superior temporal gyrus (STG), middle temporal gyrus (MTG), Heschl’s gyrus (HG), insula, anterior cingulate cortex (ACC), limbic area (combined amygdala, hippocampus, nucleus accumbens, caudate, and putamen), combined thalamus and pallidum (ThaPal), inferior frontal gyrus (IFG), inferior parietal lobule (IPL), occipital cortex, and precuneus. The selection of these ROIs was informed by prior research on the neural correlates of music, emotion, auditory, action-observation and motoric functions (Putkinen et al., 2021). ROIs implicated in visual processing such as occipital cortex and precuneus were included for null model comparison assuming that these regions would not show music-dependent responses. IFG, ThaPal and limbic region were defined using Harvard-Oxford cortical and subcortical structural atlases thresholded at 25% probability, while the rest of the ROIs were defined using the AAL2 atlas. Given music’s lateralized effects on the brain (Zatorre et al., 2002), both lateral and bilateral ROI masks were used for subsequent RSA and MVP classification analysis.

### Hyperalignment

To reduce inter-subject functional variability and enable fine-grained comparisons of representational structures across subjects, the beta values within each ROI were hyperaligned into a common representational space (Haxby et al., 2011). First, for each ROI and each subject, the data were structured into a matrix where each row represents a condition (the 20 songs for the following RSA, and the 4 emotion categories for the following MVP classification analysis), and each column a voxel (**more details in Supplementary Material**). In parallel, individual transformation matrices mapping the individual native cortical responses into the common space were computed using procrustes transformation applied to preprocessed BOLD response data obtained from the same participants during an independent fMRI task of ∼9min where they listened to a distinct and larger set of emotional music excerpts. Thus, the data used to estimate the hyperalignment transformations were separate from the data used in the present analyses. This independent data set also provides a rich and diverse set of cortical patterns that facilitates the generalization of the hyperalignment parameters to novel stimuli (Haxby et al., 2020). Each weight within the transformation matrix indicates how much the activity pattern from one voxel contributes to a dimension in the common space. Finally, for each subject, the data matrix was then multiplied by the transformation matrix, projecting the individual data into the common representational space. For the following RSA analysis, the dimensionality of the individual data was reduced into 19 components (20 song stimuli-1) using principal component analysis (PCA).

### Representational Similarity Analysis

To map brain regions where the differences in neural patterns align with the differences in bodily sensations elicited by the songs, we conducted a region-of-interest representational similarity analysis (RSA) (Kriegeskorte et al., 2008) using MATLAB’s RSA toolbox (Nili et al., 2014). First, for each subject and each ROI, the hyperaligned principal components (PC) for each song were correlated with those for every other song, constructing a representational dissimilarity matrix (RDM). Each cell in the RDM contains a dissimilarity value (1-Pearson correlation), indicating the between-song dissimilarity of the neural patterns in a given ROI. In parallel, we constructed a model RDM from the group-averaged (71 subjects) bodily sensation maps (the t-values extracted from the pixels inside the body map) by computing the dissimilaries (1-Pearson) of the BSMs across all songs. Furthermore, the extracted musical features as well as the averaged-across-subjects (68 subjects) emotion ratings for each song were used to compute two additional model RDMs for comparison. Correlations between these three model RDMs were calculated using Mantel’s test. These three similarity structures were then used to predict the similarity structure of the neural patterns. Specifically, for each ROI, the neural RDM was correlated separately with the three model RDMs using Spearman’s rank correlation. Statistical significance was assessed using the Wilcoxon signed-rank test (*p* < 0.05, FDR corrected).

### Multivariate Pattern Classification Analysis

Multivariate pattern (MVP) classification analysis was used to predict the emotion category on the basis of the brain activity elicited by the music. The data used was the hyperaligned weighted sums per emotion category computed for each subject and each ROI. Prior to the classification, features were normalized to have zero mean and unit variance. Linear support vector machine (SVM) classifier was implemented in Python, and its performance was evaluated with leave-one-subject-out cross-validation (LOSO-CV), where, for each fold, the classifier was trained on all data except a single holdout subject, and then tested on the holdout subject’s data. This process was repeated across all subjects in each ROI. Classification accuracy was defined as the proportion of correctly classified emotion categories across all subjects. For each ROI, the analysis was repeated 1000 times while randomly shuffling the category labels (aggressive, happy, sad, scary) within each subject to generate a null distribution (Pereira et al., 2009). Classification accuracy was considered significant if it exceeded the 95th percentile of the null distribution.

Pattern classification analysis was also performed using bodily sensation topographies, emotion ratings, and musical features of song to predict the emotion category. For the bodily sensations maps, the data used were the averaged-across pixels t-values throughout 6 body regions (head, trunk, shoulders, upper body (combined shoulders and trunk), arms, and legs (see **Supplementary Figure S1**. for exact body map segmentation)). Classification analysis on bodily sensations maps was additionally conducted using all the t-values extracted from the whole body as well as from each body part. Here, the t-values from the training and the testing data were separately reduced into principal components (with explained variance of at least 80%) using PCA. For the emotion ratings, the data used were the averaged-across-subjects ratings throughout the 10 rated emotions. Finally, for the musical features, the scores throughout the 20 musical features were used. SVM’s performance was tested with leave-one-song-out cross-validation.

## Results

### Bodily Sensation Topographies and Subjective Emotion Ratings Associated with Music-Induced Emotions

The bodily sensation maps revealed emotion-specific bodily activation topographies: *aggressiveness* was associated with increased sensations in the head and in the chest, *happiness* with widespread activation across the torso and extremities, *sadness* with sensations concentrated in the chest, and *fear* with heightened activation in the chest and abdomen (**Figure 1B**.).

**Figure 1.**
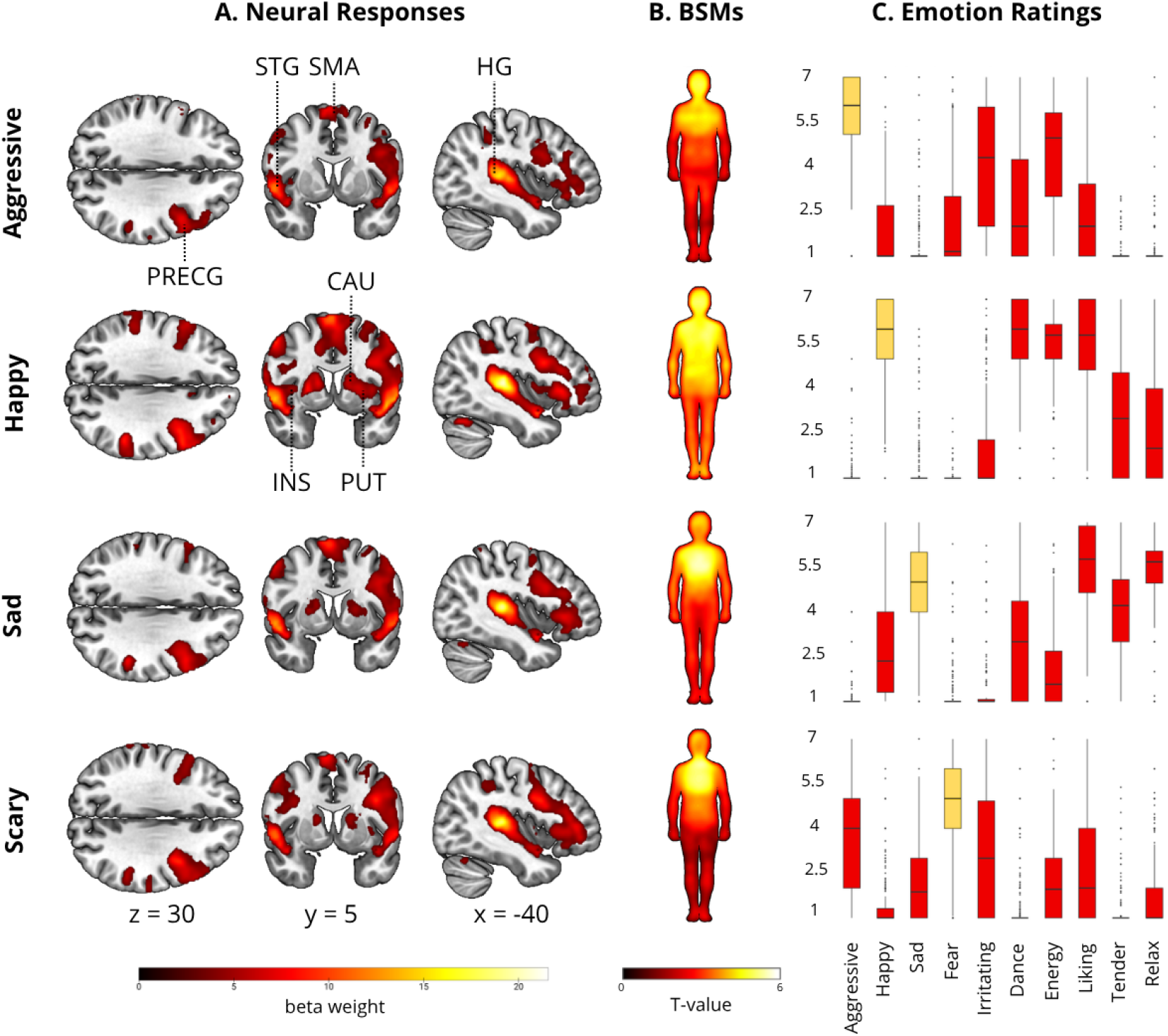
Brain responses, bodily sensation topographies and emotion ratings associated with music-induced emotions. **A**. The brain maps show regions where BOLD responses increased when listening to songs in each emotion category (aggressiveness, happiness, sadness and fear) relative to silence (*p* < 0.001 at voxel level and FDR corrected at cluster level). **B**. The body maps show regions whose activation increased when listening to songs in each category (averaged across songs within each category, *p* < 0.05 FDR corrected). **C**. The boxplots show the distribution of scores across the rated emotions for each music-induced emotion category (averaged across songs within each category for each participant). CAU caudate, INS insula, HG Helschl’s gyrus, PRECG precentral gyrus, PUT putamen, SMA supplementary motor area, STG superior temporal gyrus.

The emotion ratings were highest for emotions corresponding to the music-induced emotion category: higher ratings for aggressiveness and energy and average ratings for irritation were given to *aggressive* songs, greater ratings for happiness, danceability, liking, tenderness and relaxation were given to *happy* songs, higher ratings for sadness, liking, tenderness and relaxation were attributed to *sad* songs, and greater ratings for fear and average ratings for aggressiveness and irritation were attributed to *scary* songs (**Figure 1C**.).

The musical features were reduced into 3 PCs (**Supplementary Figure S2**.). The first PC had the highest loadings for features reflecting bright and complex timbres (spectral roll-off, spectral spread). Energy, danceability and happiness correlated the highest with this PC, and fear, sadness and relaxation the lowest. The second PC had the highest loadings for features reflecting variation in rhythmic intensity (fluctuation max), dissonance (roughness standard deviation) and in tonal profile (chroma peak standard deviation). Sadness, relaxation, tenderness and liking correlated the highest, and irritation and aggressiveness the lowest with this PC. Finally, PC3 had the highest loadings for features reflecting distorted timbre and percussive sounds (roughness mean), variation in spectral profile (spectral novelty), strong pulse (pulse clarity), and frequency of harmonic change (HCDF). Accordingly, aggressiveness, fear and irritation correlated the highest with this PC, and liking, happiness, danceability and tenderness the lowest.

### Brain Responses Associated with Music-Induced Emotions

The GLM analysis revealed significantly higher BOLD responses during music listening compared to silence across motor, auditory and emotion-related brain regions for all emotion categories (**Figure 1A**.). Specifically, activation was observed within the bilateral precentral gyrus (PRECG), supplementary motor area (SMA), superior temporal gyrus (STG), middle temporal gyrus (MTG), Heschl’s gyrus (HG), inferior frontal gyrus (IFG), and left inferior parietal lobule (IPL) across the emotion categories. *Aggressive, happy* and *sad* songs further elicited responses in left postcentral gyrus (POSCG), while *happy, sad*, and *scary* songs also activated the left putamen and bilateral caudate and supramarginal gyrus (SMG). Furthermore, *happy* and *sad* songs induced BOLD responses in bilateral pallidum as well as the left superior frontal gyrus (SFG), while *happy* and *scary* songs induced responses in bilateral insula. Finally, *happy* songs also activated bilateral thalamus and middle cingulate cortex (MCC). Overall, ROI plots of the mean beta weights per emotion category revealed higher BOLD response across all emotion categories in the STG and HG in comparison to other regions (**Figure S3**.). *Happy* songs also elicited greater activation than other emotion categories in almost all ROIs, and most notably in the STG and HG. While the mean activations overlap greatly across the emotion categories, differences, although small, can be observed between them across the ROIs, with more pronounced differences in the STG and HG. The statistical maps can be found in OSF (Rassouli et al., 2026).

### Neural Pattern Representational Similarities with Bodily Sensation Topographies, Emotion Ratings, and Musical Features

The 3 model RDMs captured the (dis)similarity structure of the songs in regard to the bodily sensations, the emotion ratings and the musical features (**Figure 2A**.). The bodily sensation maps showed weaker emotion category-wise (dis)similarity structure than the emotion ratings, although the bodily responses were still more similar within the same emotion category than between. Conversely, only the aggressive songs showed similar musical features with each other, whereas the musical features of other songs were more heterogeneous. Mantel’s test revealed a moderate but robust similarity between the bodily sensation maps and the emotion ratings’ RDMs (r = 0.326, p < 0.0001), and a weak but significant correlation between the emotion ratings and the musical features’ RDMs (r = 0.147, p < 0.05).

**Figure 2.**
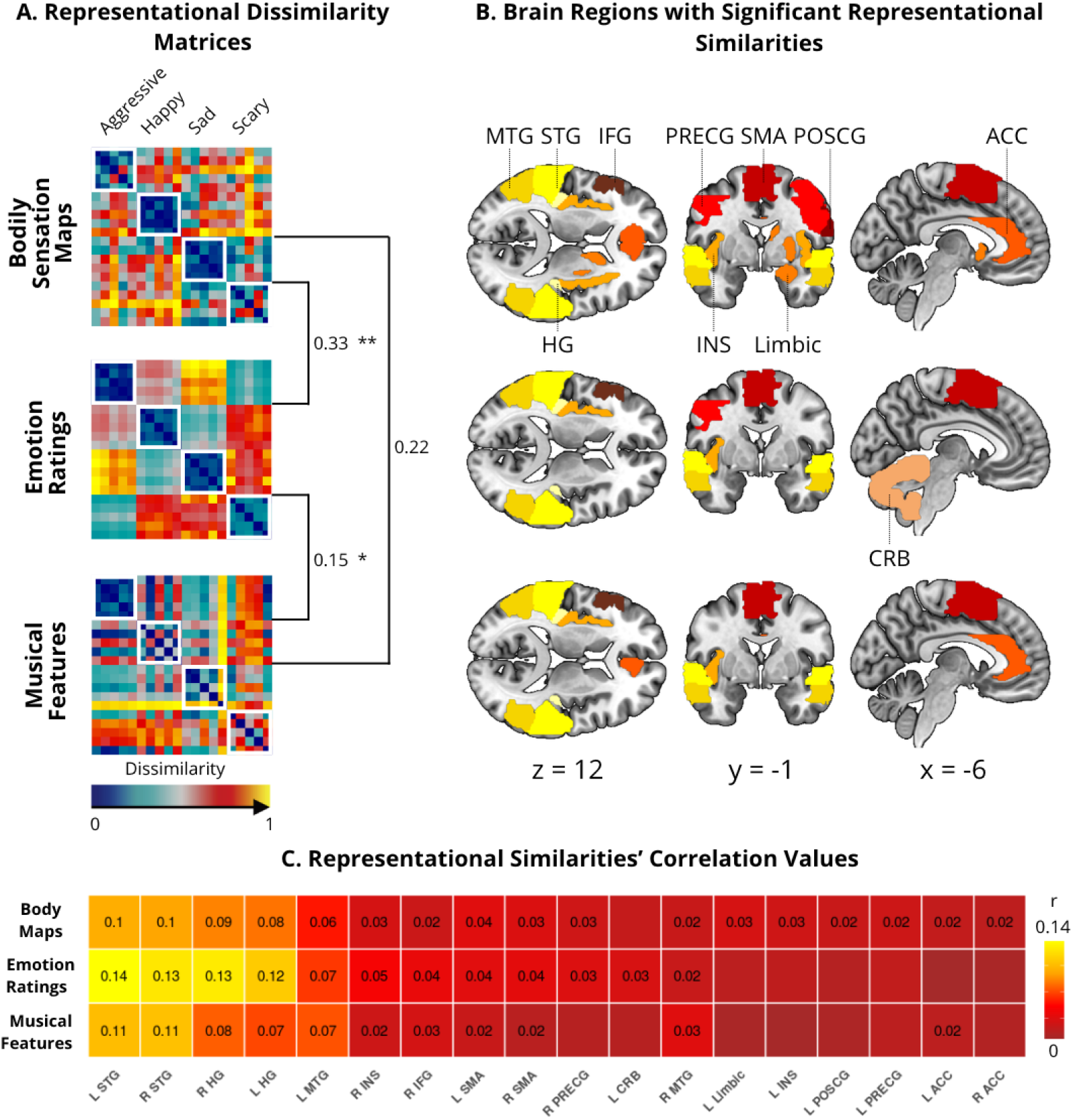
Representational Similarities of Neural Patterns with Bodily Sensations, Emotion Ratings and Musical Features. **A**. The representational dissimilarity matrices (RDMs) of the group-averaged music-evoked bodily sensations and emotion ratings, as well as song-wise musical features show which songs are associated with similar/dissimilar bodily sensations, ratings, and musical features. Dissimilarity is shown as 1-Pearson’s correlation value, rank-transformed and scaled [0,1] with 0 describing perfect similarity and 1 dissimilarity. The white squares highlight the emotion category boundaries. The double-headed arrows indicate the correlations between these model RDMs (* for *p* < 0.05, ** for *p* < 0.0001). **B**. The brain figures show the ROIs whose neural patterns’ representational geometries are significantly correlated with the representational geometries of the bodily sensations topographies, the emotion ratings, and the musical features. **C**. The heatmap indicates the spearman rank’s correlation values, and the significant correlations (*p* < 0.05 FDR corrected) are highlighted with the correlation value r. ACC anterior cingulate gyrus, CRB cerebellum, IFG inferior frontal gyrus, Limbic limbic region (combining amygdala, hippocampus, nucleus accumbens, caudate, and putamen), MTG middle temporal gyrus, POSCG postcentral gyrus.

Altogether, RSA revealed significant correspondences between the representational geometries of neural activity patterns and the three model RDMs (**Figure 2B. and 2C**.). Interestingly, similarities with the bodily sensation map’s model were most consistently observed across the ROIs. Specifically, somatomotor areas such as the bilateral PRECG and SMA, the left POSCG and the right IFG revealed significant similarities with the bodily sensation maps’ geometries, while only the bilateral SMA, right PRECG and IFG, and left cerebellum showed significant similarities with the emotion ratings’, and only the bilateral SMA and the right IFG with the musical features’ geometries. Additionally, bilateral ACC, left limbic regions and regions of the canonical emotion network such as the bilateral insula revealed significant similarities with the bodily sensation map’s model. Musical features’ and emotion ratings’ representational structures also showed significant similarities with the right insula, and musical features’ model with the left ACC as well. Finally, the auditory cortex (bilateral STG, MTG and HG) showed significant similarities with all model’s representational dissimilarity structures. Taken together, bilateral SMA, STG, MTG and HG, and the right insula and IFG showed significant correlation with all three model’s RDM. In general, strongest correlations were observed within the auditory cortex. Occipital cortex, precuneus, ThaPal and IPL ROIs did not reveal significant similarities with any of the three models’ representational geometries. The analyses were performed using bilateral ROIs as well, revealing similar although less specific results as analyses computed on lateral ROIs, and were thus not discussed here.

### Emotion Classification with Multivariate Pattern Classification Analysis

The ROI-level MVP classification analysis revealed that the classification accuracy for all four emotions was significant for the auditory cortex (STG and HG with overall classification accuracy of 50% and 39% respectively) (**Figure 3**.). While the overall classification accuracy significantly exceeded the chance level for PRECG (35%), insula (31%), limbic regions (31%), and SMA (30%), none of these ROIs reliably discriminated between all four emotions. Furthermore, their classification accuracies were considerably lower than that of the STG and HG. The analyses were performed separately for the left and right hemispheres as well, yet no noteworthy hemispheric differences were observed and the classification accuracies for the bilateral analyses tended to be higher than for the lateralized ones.

**Figure 3.**
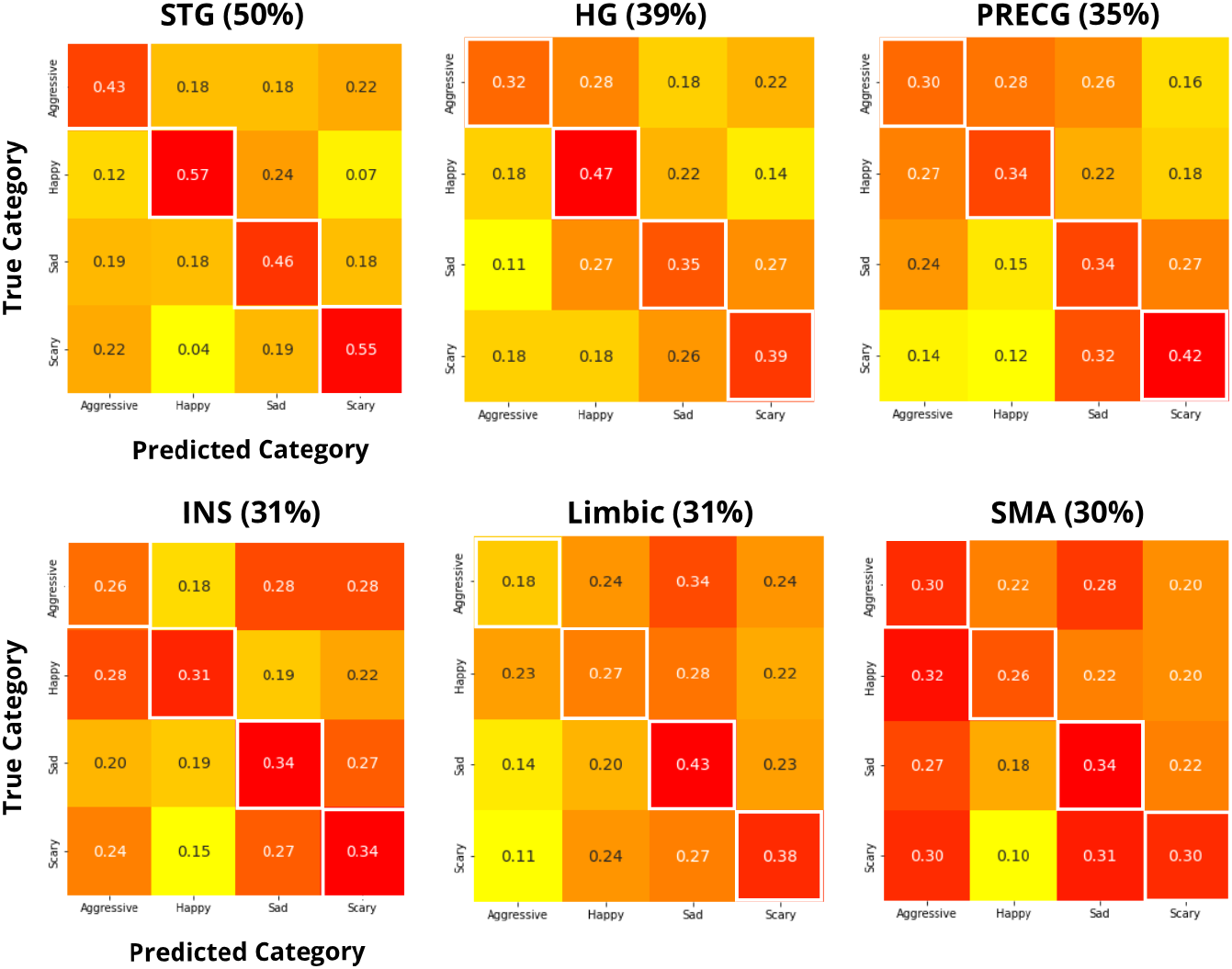
MVP classification analysis confusion matrices for the ROIs with significant overall classification accuracy. The numbers indicate the proportion of correct and false predictions for each emotion category with true positives shown on the diagonal and misclassifications off-diagonal.

In parallel, the prediction of emotion categories using body maps, emotion ratings and musical features were all significantly above chance level with overall classification accuracies of 85%, 100% and 80%, respectively (**Supplementary Figure S4**.).

## Discussion

Our main finding was that representational similarities between emotion-specific music-evoked bodily sensation topographies and neural activation patterns are shared in motor and somatosensory cortices. Interestingly, representational correspondence was also found across neural networks beyond the somatomotor regions, such as auditory cortex, limbic and interoceptive brain regions, suggesting that the embodiment of musical emotions is not restricted to sensorimotor representations, but involves distributed neural systems supporting auditory, affective, and internal bodily processing. These results were observed in the absence of notable differences in GLM-based analysis of the emotion-specific neural responses to emotional music. These results indicate that distinct music-induced emotions trigger distinct bodily sensations, and this discrete organization is reflected in the neural responses.

### Music-induced subjective bodily sensations

Distinct music-induced emotions elicited distinct bodily sensations which could be accurately classified from each other (**Figure 1B. and S4**.). This accords with previous findings on emotion-specific bodily responses to music (Putkinen et al., 2024). Other studies have also used bodily sensation maps to investigate embodied feelings when listening to music. For instance, distinct patterns of embodied feelings for different components of musical groove have been revealed, where wanting to move was associated with more sensations in the extremities, and pleasure was more associated with sensations in the chest and abdomen (Witek et al., 2025). Even relatively low-level manipulations of musical structure may produce differentiated bodily sensations: a study on bodily sensations related to uncertainty and surprise in musical chord progression suggested that music-induced sensations in the chest may be related to the embodied affective dimensions of musical predictive processing (Daikoku et al., 2024). Altogether, our results along with previous findings indicate that music-induced emotional experiences are systematically embodied so that different aspects of these experiences induce distinct spatial patterns of bodily sensations.

### Neural representation of embodied music-induced emotions in the somatomotor circuits

RSA analysis revealed that information for embodied music-evoked emotional experiences was represented in brain regions associated with motor and somatosensory function (pre- and postcentral gyri, supplementary motor area, and inferior frontal gyrus) as well as emotion, interoceptive (insula, anterior cingulate cortex, and limbic region) and auditory processing (superior and middle temporal gyri, and Heschl’s gyri) (**Figure 2**.). The pre- and postcentral gyri are of particular interest given that they contain somatotopically organized primary motor and somatosensory cortices suggesting a link between the topography of music-induced bodily sensations and multivariate activation patterns in these sensorimotor regions.

A large body of literature has indicated motor system activation during music listening with consistent activation across studies in bilateral premotor cortex, right primary motor cortex, and left cerebellum (for a meta-analysis see Gordon, Cobb & Balasubramaniam, 2018), and reliable discrimination between music-induced emotions in the primary motor cortex (Putkinen et al., 2021). Such motor system activation does not merely reflect overt movement or movement preparation: for instance, motor regions such as the mid premotor cortex, the SMA and the cerebellum are similarly recruited in the absence of any anticipated movements as in presence of movement anticipation during music listening (Chen et al., 2008). Theoretical models such as the Action Simulation for Auditory Prediction (ASAP, Patel and Iversen, 2014), suggest the role of auditory-motor interactions in generating temporal prediction during music listening providing anticipatory representations about the upcoming sounds. The confirmation or disconfirmation of these expectations, in turn, have been linked with music-induced pleasure and desire to move. Consistent with this view, musical rhythms inducing higher subjective pleasure and desire to move elicit stronger neural responses in regions implicated in beat-based timing (such as supplementary motor area, putamen, parietal and prefrontal cortices) and reward processes (such as nucleus accumbens, caudate and medial orbitofrontal cortex). These findings motivated a model in which reward processing and motor processes driving rhythmic expectations influence the sensation of groove (Matthews & Witek, 2020). However, while these accounts of motor activation during music-listening emphasize prediction and sensorimotor simulation, they do not reveal emotion-specific linkage and correspondence between bodily and neural responses to music. Our findings go beyond these results by showing that activity patterns in motor and somatosensory regions share a representational structure with subjective bodily sensations evoked by music. Viewed within the predictive framework, our findings raise the possibility that emotionally distinct musical excerpts may evoke distinct anticipatory states, which contribute to subjective bodily sensations and are reflected in the representational structure of motor and somatosensory regions.

### Distributed representations of musical bodily sensations, emotions and acoustic features

Our results further suggest that information related to musical features, subjective emotional experiences, and music-induced bodily sensations is represented across distributed cortical systems extending beyond somatomotor regions (**Figure 2**.). Auditory cortical regions showed particularly robust representational similarity structure across bodily sensations, musical features, and subjective emotion dimensions, supporting the view that auditory cortex contributes not only to low-level acoustic analysis but also to higher-order affective and experiential aspects of music perception. Interestingly, bodily sensation models showed even broader and more consistent representational structure across regions than models based solely on felt emotion ratings or musical features. This pattern was also observed in limbic and emotion-related regions, where only the bodily sensation models reached significance. These findings are compatible with embodied and interoceptive accounts of musical emotion, which propose that music-evoked emotional experiences are closely linked to representations of bodily states and action tendencies. Altogether, these results reveal that even regions without direct sensorimotor functions encode information about music-induced bodily sensations. At the same time, information related to subjective emotions and musical features was also represented in somatomotor, interoceptive, and emotion-related regions, suggesting that these systems participate more broadly in the representation of affective and experiential aspects of music listening. Given the substantial interdependence between musical structure, bodily sensations, and emotional experiences, the present findings are more consistent with distributed and overlapping representations of musical, bodily, and emotional information than with sharply segregated functional systems. In line with this, here, musical features are consistently linked to particular emotions, which in turn correspond to specific bodily sensation patterns. This is further evidenced by musical feature similarity reliably predicting emotional similarity, which in turn serves as a strong predictor of bodily sensation similarity. Although this pattern does not establish a causal pathway, it is consistent with subjective emotional experience mediating the relationship between musical features and bodily sensations.

### Univariate and multivariate responses to music are not emotion category-specific beyond the auditory cortex

GLM analysis reveals that emotionally engaging music activates brain regions previously reported to be involved in emotional music processing (Putkinen et al., 2021; Koelsch et al., 2018; Koelsch et al., 2014) (**Figure 1A**.). The resulting neural activity patterns, however, overlap considerably across the emotion categories (**Figure S3**.), highlighting the limits of GLM analysis and, thus the need for multivariate approaches in discriminating the neural representations of distinct emotions and bodily sensation topographies.

Here, the MVP classification analysis could accurately classify emotion-specific responses in the auditory cortex, further validating previous findings (Putkinen et al., 2021) and highlighting its key role in emotional musical stimuli processing (**Figure 3**.). Significant emotion category classification was also observed in the precentral gyri, the supplementary motor area and the limbic regions, in accordance with Putkinen’s (2021) findings, as well as in the insula, albeit with lower discrimination between the emotion categories than within the auditory cortex.

While the bodily sensation maps, the GLM analysis and the multivariate pattern classification analysis validate previous findings, the RSA provides novel evidence for the neural basis of embodied musical emotions by revealing that neural representational geometries closely mirror the geometries of the bodily sensation maps in a distributed network comprising of the motor, somatosensory, interoceptive, limbic and auditory regions, hence suggesting that embodied experiences of musical emotions are reflected in shared representational structures across the brain and the body. A previous study employing RSA failed to find statistical similarity between neural activity patterns and bodily sensation topographies during emotion generation by recall of emotional events (Giraud et al., 2024). In contrast, we investigated real-time music-induced emotions in a larger sample and used hyperalignment to align the individual brain responses. Our results demonstrate the utility of RSA as an analytical tool to understand brain-body interaction in human emotions and encourage its use for further research on the embodiment of other types of emotions.

## Limitations

Our results support the idea that the somatomotor representations play a central role in music-induced emotions. However, it is important to note that actual bodily processes cannot be inferred from subjective bodily sensations. While these subjective bodily sensations may arise from skeletomuscular and ANS activity, they may also emerge from internal models of bodily reactions reenacting physiological responses typically elicited during emotional states (Wöllner, 2025; Craig, 2008; Damasio, 2004). This might be reflected in the reported nature of the subjective bodily sensations where the participants may feel emotional states at the physical level without actually receiving any feedback from the body. Future research clarifying whether actual bodily processes are involved, using total-body positron emission tomography (PET) imaging for instance, would inform us greatly on brain-body interaction. It is also important to note that emotions can be experienced without peripheral sensations. Indeed, emotional responses induced by music may still be shaped by top-down interoception and awareness processes (Wöllner, 2025; Craig, 2008; Damasio, 2004).

## Conclusion

Bodily sensations induced by listening to emotionally engaging music are represented in a distributed network of neural systems. Specifically, the representational geometries of the distinct music-induced bodily sensations closely match the representational geometries of the evoked neural activation patterns within the motor, somatosensory, limbic, interoceptive and auditory brain regions. These results demonstrate neural correspondence to bodily sensations during emotional experiences. We believe that future research on brain-body interaction during emotional states would further elucidate the bodily processes involved in the subjective experience of emotions.

## Data, Materials, and Software Availability

Code and statistical brain maps have been deposited in OSF (Rassouli et al., 2026).

## Author contributions

**Maya Rassouli**: Formal analysis, Investigation, Data curation, Writing - Original draft, Writing - Review and editing, Visualization. **Viljami Salmela**: Resources, Writing - Review and editing. **Kerttu Seppälä**: Investigation. **Harri Harju**: Investigation. **Lauri Nummenmaa**: Conceptualization, Writing - Review and editing. **Vesa Putkinen**: Conceptualization, Writing - Review and editing, Supervision, Funding acquisition.

## Acknowledgments

We sincerely thank Eveliina Rantakylä, Anssi Rinne, and Tuomo Ouvinen for their contribution to participant recruitment and data collection.

Supported by the Signe & Ane Gyllenbergs Foundation grant to MR, by the European Research Council (ERC-ADV #101141656) and the Jane & Aatos Erkko Foundation to LN, as well as Academy of Finland to VP.

## Supplementary Material

**Table S1.**
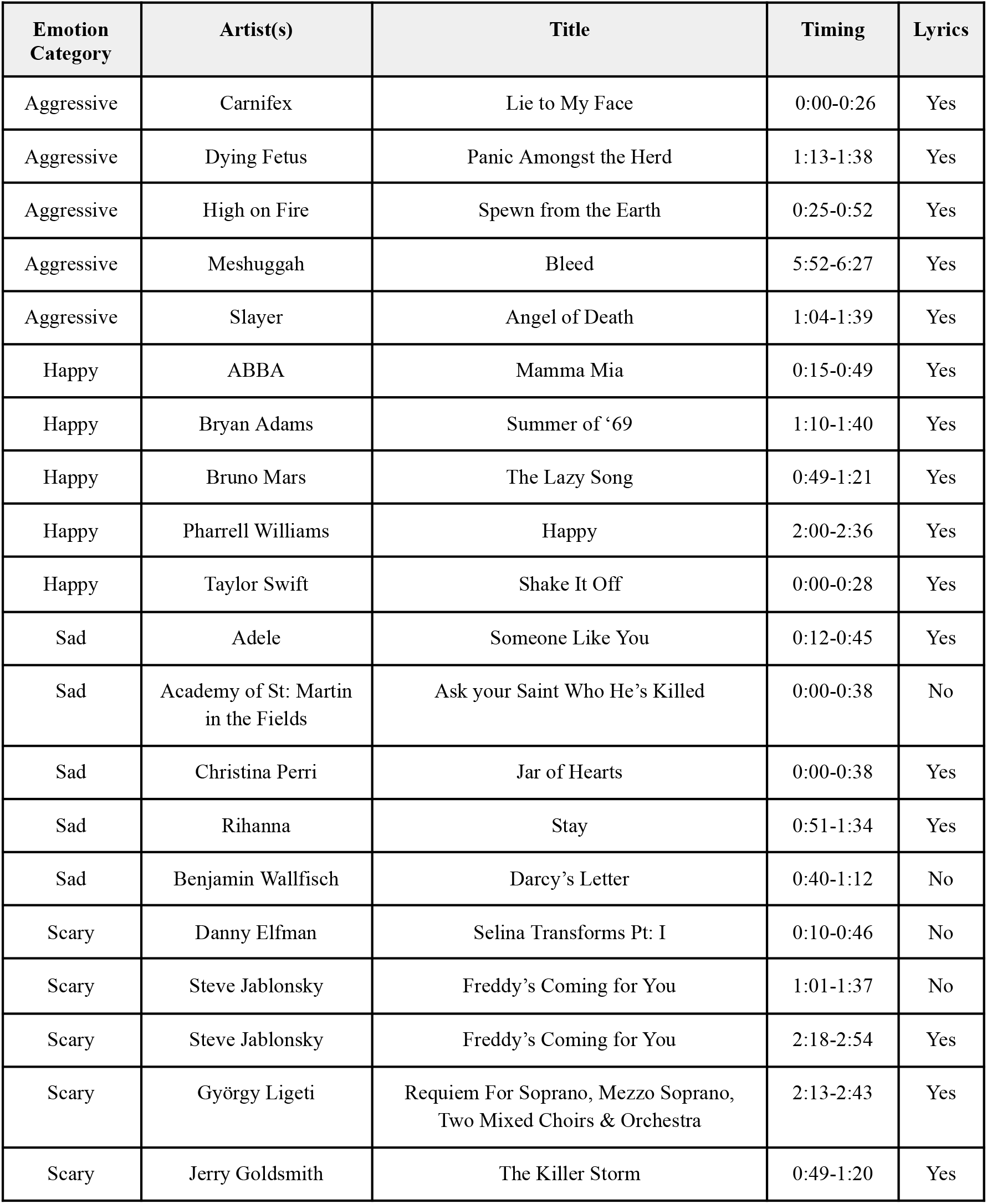
List of the Songs.

| Emotion Category | Artist(s) | Title | Timing | Lyrics |
| --- | --- | --- | --- | --- |
| Aggressive | Carnifex | Lie to My Face | 0:00-0:26 | Yes |
| Aggressive | Dying Fetus | Panic Amongst the Herd | 1:13-1:38 | Yes |
| Aggressive | High on Fire | Spewn from the Earth | 0:25-0:52 | Yes |
| Aggressive | Meshuggah | Bleed | 5:52-6:27 | Yes |
| Aggressive | Slayer | Angel of Death | 1:04-1:39 | Yes |
| Happy | ABBA | Mamma Mia | 0:15-0:49 | Yes |
| Happy | Bryan Adams | Summer of '69 | 1:10-1:40 | Yes |
| Happy | Bruno Mars | The Lazy Song | 0:49-1:21 | Yes |
| Happy | Pharrell Williams | Happy | 2:00-2:36 | Yes |
| Happy | Taylor Swift | Shake It Off | 0:00-0:28 | Yes |
| Sad | Adele | Someone Like You | 0:12-0:45 | Yes |
| Sad | Academy of St: Martin in the Fields | Ask your Saint Who He's Killed | 0:00-0:38 | No |
| Sad | Christina Perri | Jar of Hearts | 0:00-0:38 | Yes |
| Sad | Rihanna | Stay | 0:51-1:34 | Yes |
| Sad | Benjamin Wallfisch | Darcy's Letter | 0:40-1:12 | No |
| Scary | Danny Elfman | Selina Transforms Pt: I | 0:10-0:46 | No |
| Scary | Steve Jablonsky | Freddy's Coming for You | 1:01-1:37 | No |
| Scary | Steve Jablonsky | Freddy's Coming for You | 2:18-2:54 | Yes |
| Scary | György Ligeti | Requiem For Soprano, Mezzo Soprano, Two Mixed Choirs & Orchestra | 2:13-2:43 | Yes |
| Scary | Jerry Goldsmith | The Killer Storm | 0:49-1:20 | Yes |

**Table S2.**
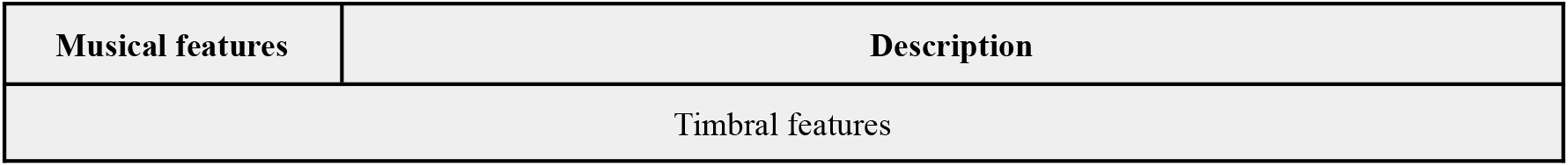
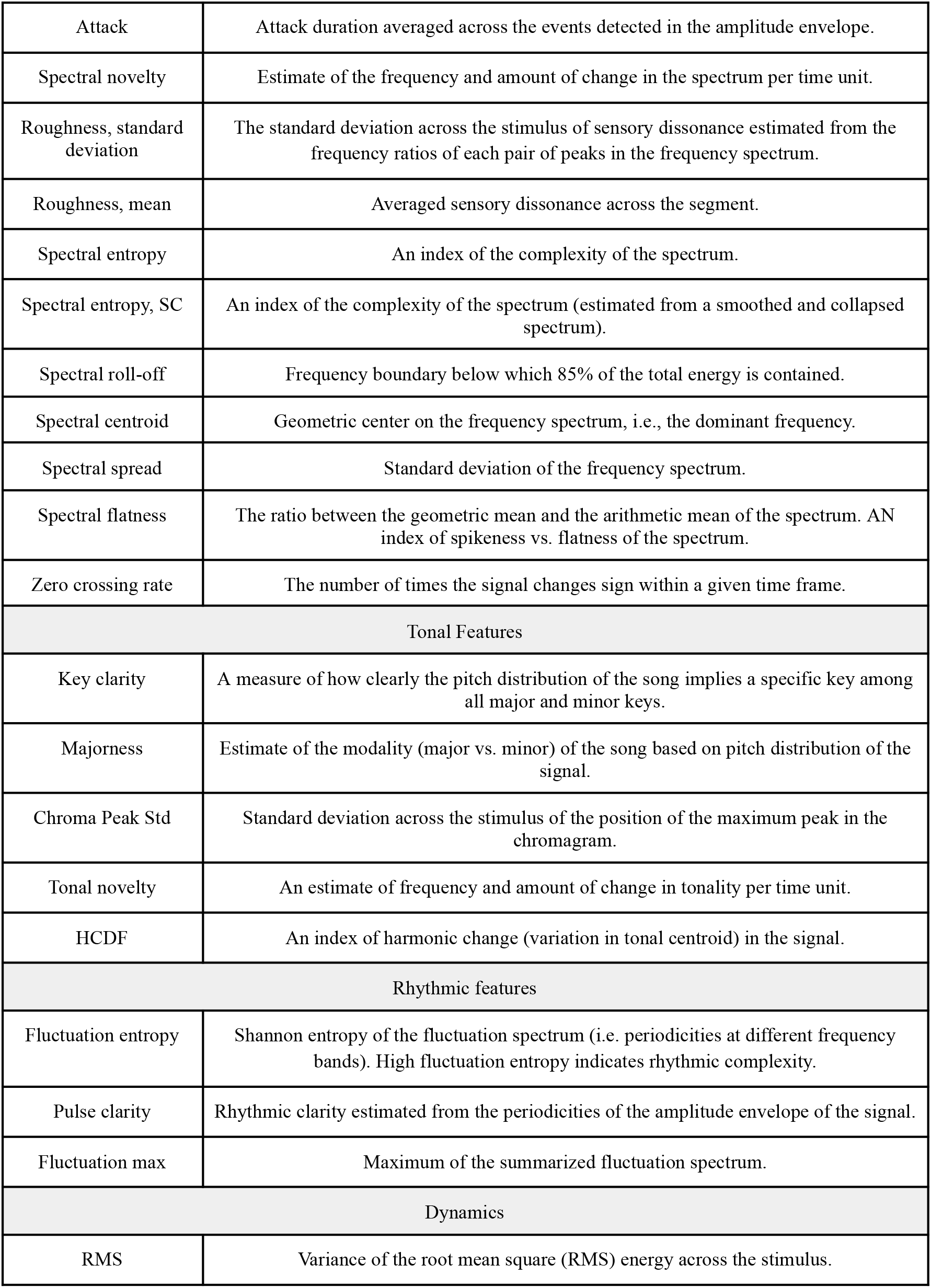
Musical Features.

| Musical features | Description |
| --- | --- |
| Timbral features |  |
| Attack | Attack duration averaged across the events detected in the amplitude envelope. |
| Spectral novelty | Estimate of the frequency and amount of change in the spectrum per time unit. |
| Roughness, standard deviation | The standard deviation across the stimulus of sensory dissonance estimated from the frequency ratios of each pair of peaks in the frequency spectrum. |
| Roughness, mean | Averaged sensory dissonance across the segment. |
| Spectral entropy | An index of the complexity of the spectrum. |
| Spectral entropy, SC | An index of the complexity of the spectrum (estimated from a smoothed and collapsed spectrum). |
| Spectral roll-off | Frequency boundary below which 85% of the total energy is contained. |
| Spectral centroid | Geometric center on the frequency spectrum, i.e., the dominant frequency. |
| Spectral spread | Standard deviation of the frequency spectrum. |
| Spectral flatness | The ratio between the geometric mean and the arithmetic mean of the spectrum. AN index of spikeness vs. flatness of the spectrum. |
| Zero crossing rate | The number of times the signal changes sign within a given time frame. |
| Tonal Features |  |
| Key clarity | A measure of how clearly the pitch distribution of the song implies a specific key among all major and minor keys. |
| Majorness | Estimate of the modality (major vs. minor) of the song based on pitch distribution of the signal. |
| Chroma Peak Std | Standard deviation across the stimulus of the position of the maximum peak in the chromagram. |
| Tonal novelty | An estimate of frequency and amount of change in tonality per time unit. |
| HCDF | An index of harmonic change (variation in tonal centroid) in the signal. |
| Rhythmic features |  |
| Fluctuation entropy | Shannon entropy of the fluctuation spectrum (i.e. periodicities at different frequency bands). High fluctuation entropy indicates rhythmic complexity. |
| Pulse clarity | Rhythmic clarity estimated from the periodicities of the amplitude envelope of the signal. |
| Fluctuation max | Maximum of the summarized fluctuation spectrum. |
| Dynamics |  |
| RMS | Variance of the root mean square (RMS) energy across the stimulus. |

## Supplementary Methods

### Hyperalignment

For lateralized ROI masks exceeding 2000 voxels, the best 2000 voxels (the voxels showing the highest BOLD responses) were selected from the ROI-masked beta maps of each condition and each individual. For ROI masks encompassing both hemispheres exceeding 4000 voxels, the best 4000 voxels were selected. In case the number of voxels within the ROI is lower than 2000 or 4000 voxels accordingly, all the voxels within that ROI were selected. Lateralized ROI masks were used for RSA, while masks encompassing both hemispheres were used for MVP classification analysis.

## Supplementary Figures

**Figure S1.**
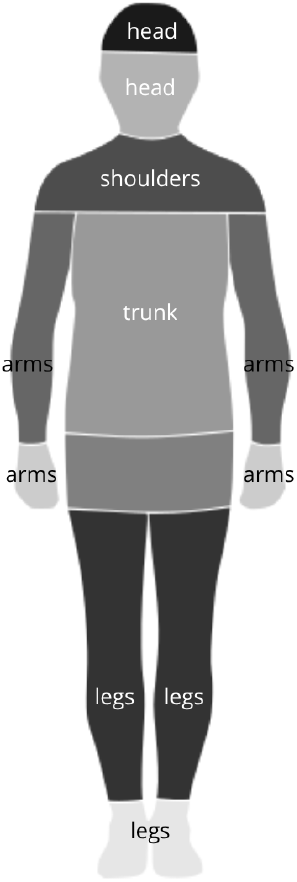
Body maps segmentation.

**Figure S2.**
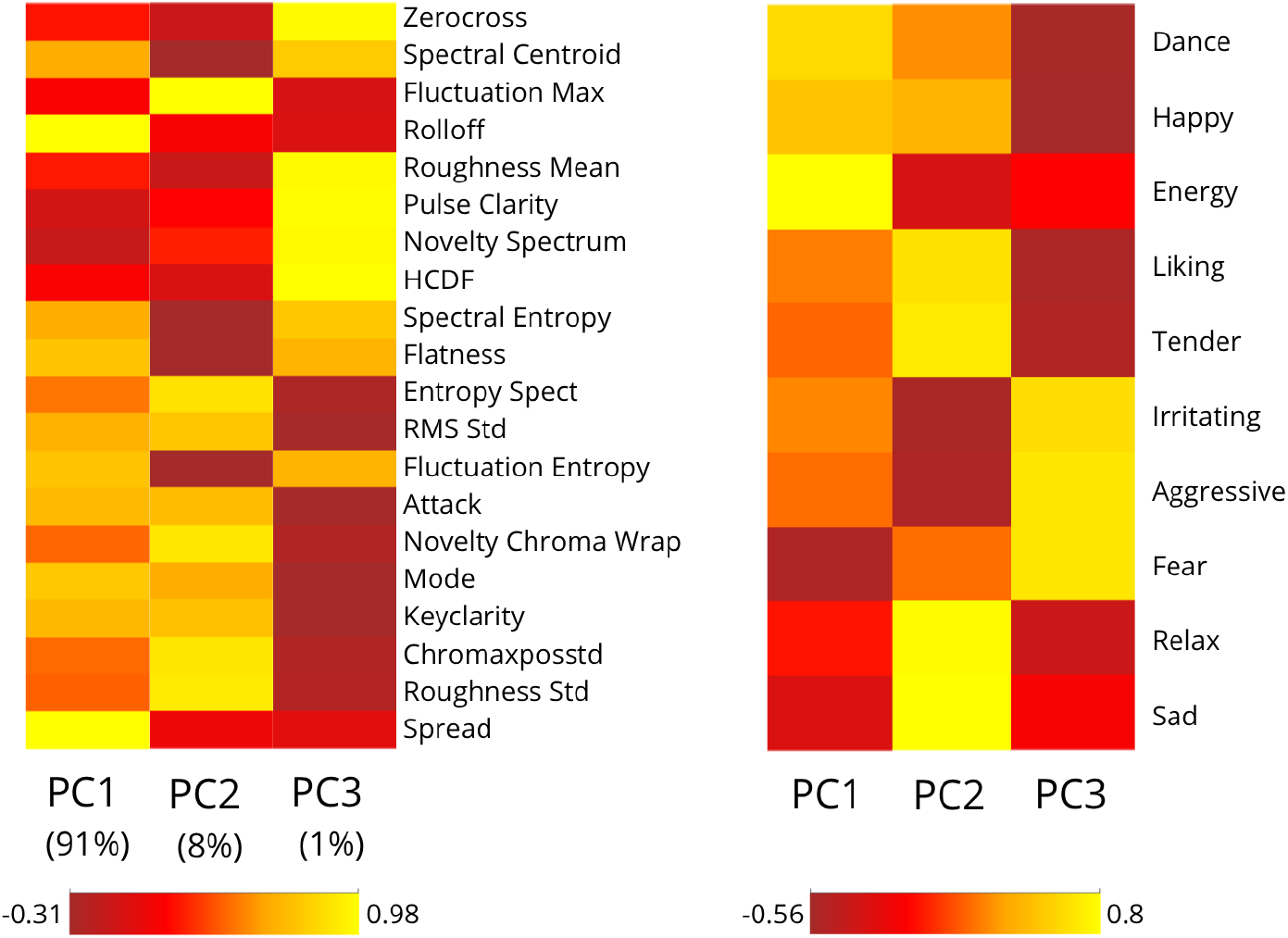
Musical Features’ principal components’ coefficients and explained variance, and their correlation with Emotion Ratings.

**Figure S3.**
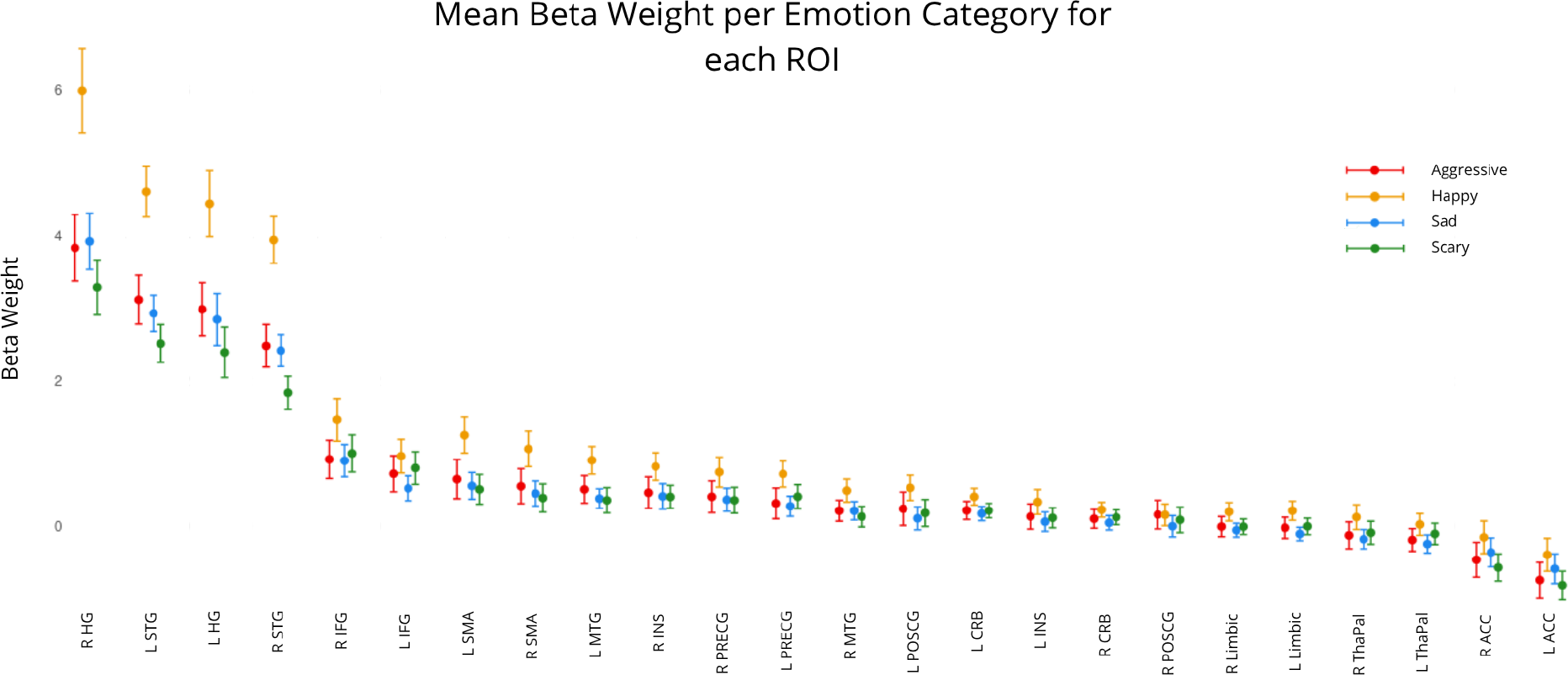
Mean beta weight per emotion category for each ROI. The mean beta weight and the 95% confidence interval for each emotion category was computed across subjects and plotted for distinct regions-of-interests.

**Figure S4.**
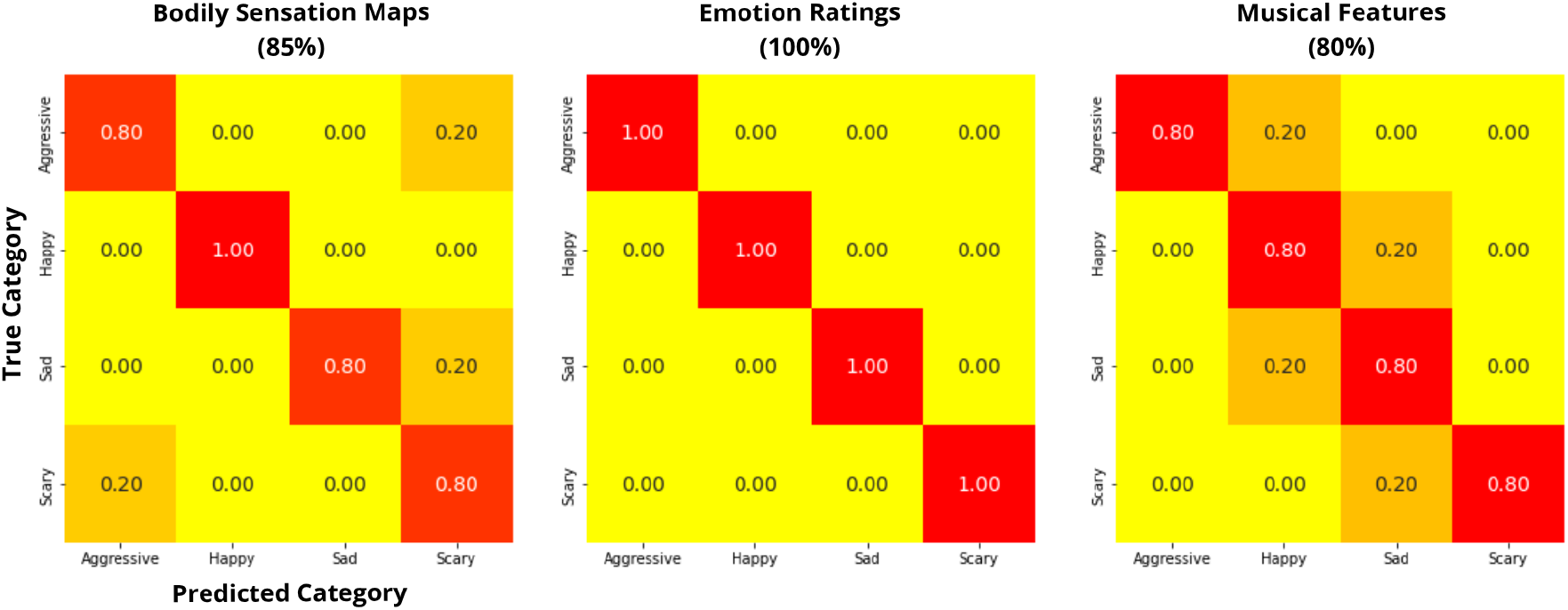
MVP classification analysis confusion matrices and overall classification accuracies based on Bodily Sensation Maps ROI-wise mean t-values, Emotion Ratings mean ratings and Musical Features.

**Figure S5.**
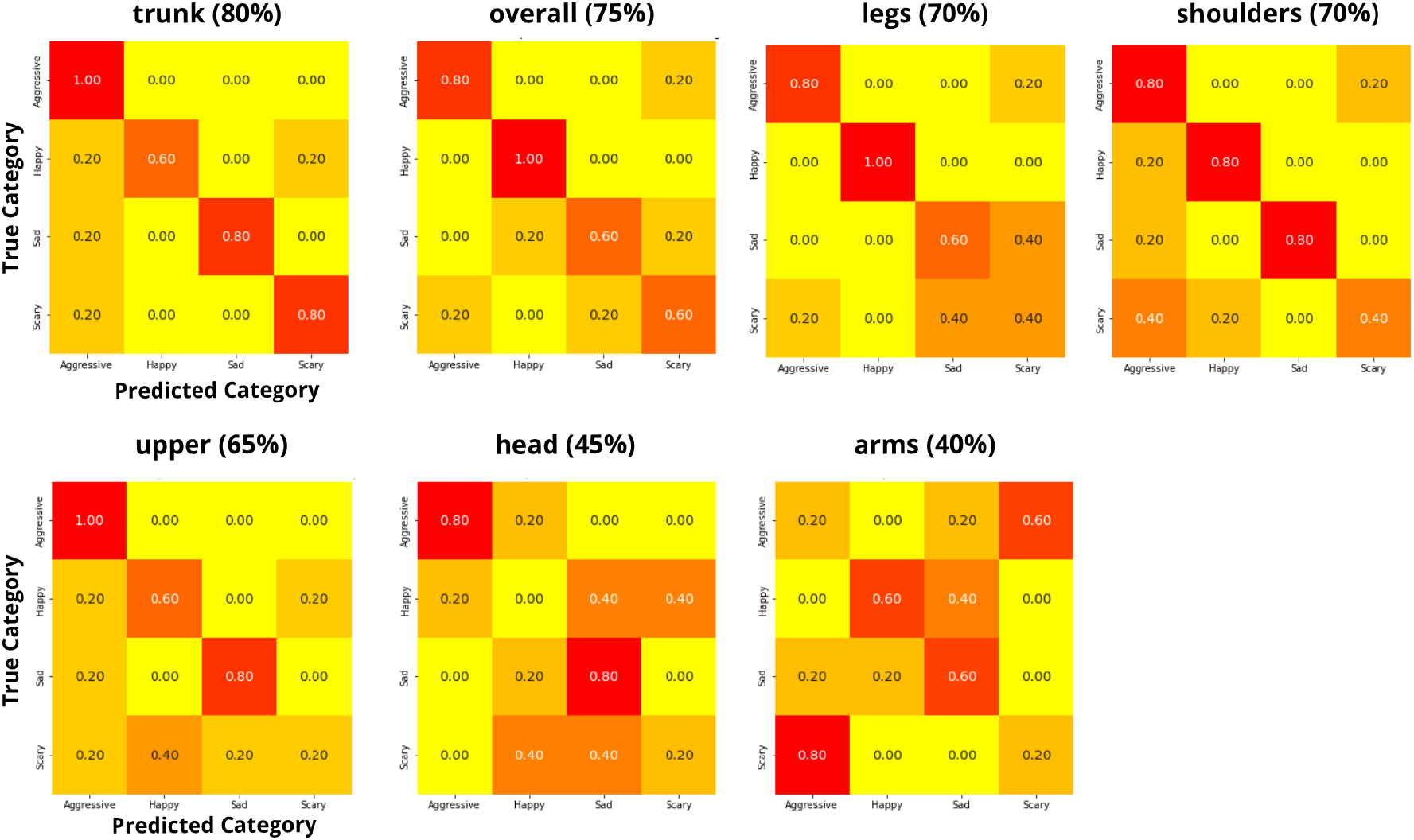
MVP classification analysis confusion matrices and overall classification accuracies based on Bodily Sensation Maps t-values.

